# Selectivity of Guanine Nucleotide Exchange Factor-mediated Cdc42 activation in primary human endothelial cells

**DOI:** 10.1101/541219

**Authors:** Nathalie R. Reinhard, Sanne van der Niet, Anna Chertkova, Marten Postma, Peter L. Hordijk, Theodorus W.J. Gadella, Joachim Goedhart

## Abstract

The Rho GTPase family is involved in actin dynamics and regulates the barrier function of the endothelium. One of the main barrier-promoting Rho GTPases is Cdc42, also known as cell division control protein 42 homolog. Currently, regulation of Cdc42-based signaling networks in endothelial cells (ECs) lack molecular details. To examine these, we focused on a subset of 15 Rho guanine nucleotide exchange factors (GEFs), which are expressed in the endothelium. By performing single cell FRET measurements with Rho GTPase biosensors in primary human ECs, we monitored GEF efficiency towards Cdc42 and Rac1. A new, single cell-based analysis was developed and used to enable the quantitative comparison of cellular activities of the full-length GEFs. Our data reveal a specific GEF dependent activation profile, with most efficient Cdc42 activation induced by PLEKHG2, FGD1, PLEKHG1 and pRex1 and the highest selectivity for FGD1. Additionally, we generated truncated GEF constructs that comprise only the catalytic dbl homology (DH) domain or together with the adjacent pleckstrin homology domain (DHPH). The DH domain by itself did not activate Cdc42, whereas the DHPH domain of ITSN1, ITSN2 and PLEKHG1 showed activity towards Cdc42. Together, our study characterized endothelial GEFs that may activate Cdc42, which will be of great value for the field of vascular biology.

Graphical Abstract

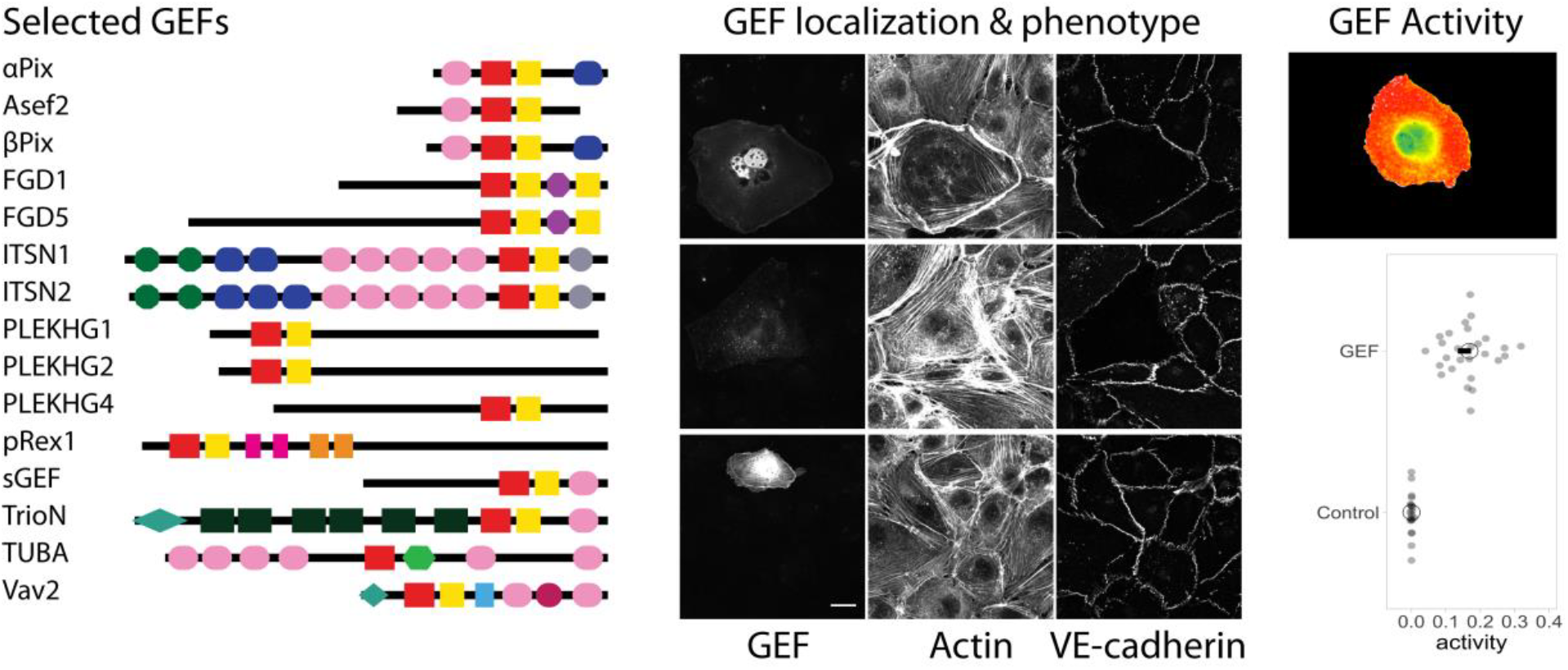

## Introduction

The Rho family of small GTPases belongs to the superfamily of Ras GTPases. Approximately 20 Rho GTPases have been identified, some of which show >85% homology and functional redundancy^1,2^. Via differential actin cytoskeleton remodeling, Rho GTPases regulate a range of cellular responses, including cell adhesion, -migration and -polarity^3^. In the endothelium, these processes form the basis of vascular homeostasis and dynamic regulation of endothelial barrier function. Consequently, Rho GTPases are key molecular components in EC biology.

Rho GTPases act as molecular switches, and their activation and downstream signaling is regulated by three groups of proteins^4^. While Rho guanine nucleotide exchange factors (GEFs) mediate the Rho GTPase GDP-GTP exchange, leading to an activated GTPase, the intrinsic GTP hydrolysis, i.e. inactivation, is stimulated by Rho GTPase activating proteins (GAPs). Additionally, members of the Rho guanine dissociation inhibitors (GDI) family, sequester inactive Rho GTPases in the cytoplasm, securing a large pool of Rho GTPases available for rapid cellular responses to external cues. Roughly 80 GEFs, 70 GAPs and 3 GDIs have been identified, greatly outnumbering the ∼20 Rho GTPases. Consequently, the regulation of Rho GTPase activity, both in time and place, is highly complex^3,4^.

GEFs comprise the largest group of Rho GTPase regulators, consisting of two families; 11 dedicator of cytokinesis (Dock) - and 74 diffuse B-cell lymphoma (Dbl) proteins^5–13^. The Dbl family has been repeatedly linked to Rho activation. Members of this family contain a Dbl-homology (DH) domain of around 170-190 residues, that regulates the Rho GTPase GDP-GTP exchange^10,13^. In addition, the majority of DH-containing Dbl family members encode a pleckstrin homology (PH) domain of approximately 120 residues. PH domains can contribute to GEF autoinhibition, activity regulation, subcellular localization, phospholipid binding and to the scaffolding of related signaling proteins^4,10,14^. However, these functions are relatively unexplored and in general PH-domain specific functions are poorly documented.

Although GEF-Rho GTPase interactions are critical determinants in Rho GTPase signaling networks, molecular details are still missing. Additionally, current studies are mostly based on studies with isolated components or pull-down experiments on lysed cells. The *in vitro* data provide mechanistic details^13^, but lack physiological relevance due to the absence of cellular context. Here, we specifically focused on GEF-Cdc42 interactions in the cellular environment, with special emphasis on the endothelium. We selected a subset of endothelial GEFs and performed fluorescence resonance energy transfer (FRET) measurements to measure Cdc42 activation in live primary HUVECs (human umbilical vein endothelial cells). This single-cell FRET-based approach identifies critical activators of Cdc42 in the endothelium.

## Results

### Cdc42 GEF selection in human ECs

Over the past years the Rho GTPase Cdc42 has been identified as a key regulator in endothelial barrier control^15–18^. Endothelial barrier function is dependent on tight orchestration of complex, interacting signaling pathways, of which detailed information is still missing. To explore Cdc42-mediated signaling networks in ECs, we here tested a series of potential Cdc42 GEFs for their selectivity and effectivity in activating Cdc42.

Potential Cdc42 GEFs were selected, based on (i) expression level analysis; (ii) catalytic Rho GEF activity (mostly based on in vitro studies); and (iii) reported effects on (barrier) function in ECs^13,19–21^. Following these criteria, 15 GEFs were identified to be of interest; these include α-Pix, Asef2, β-Pix, FGD1, FGD5, ITSN1, ITSN2, PLEKHG1, PLEKHG2, PLEKHG4, pRex1, sGEF, TrioN, TUBA and Vav2. Except for TUBA (lacking PH domain), these GEFs all contain a DH and a PH domain, positioning them as members of the Dbl GEF family. A schematic overview of the domain organization within these GEFs was obtained using the SMART (Simple Modular Architecture Research Tool) database^22^, and is illustrated in Figure 1.

**Figure 1.**
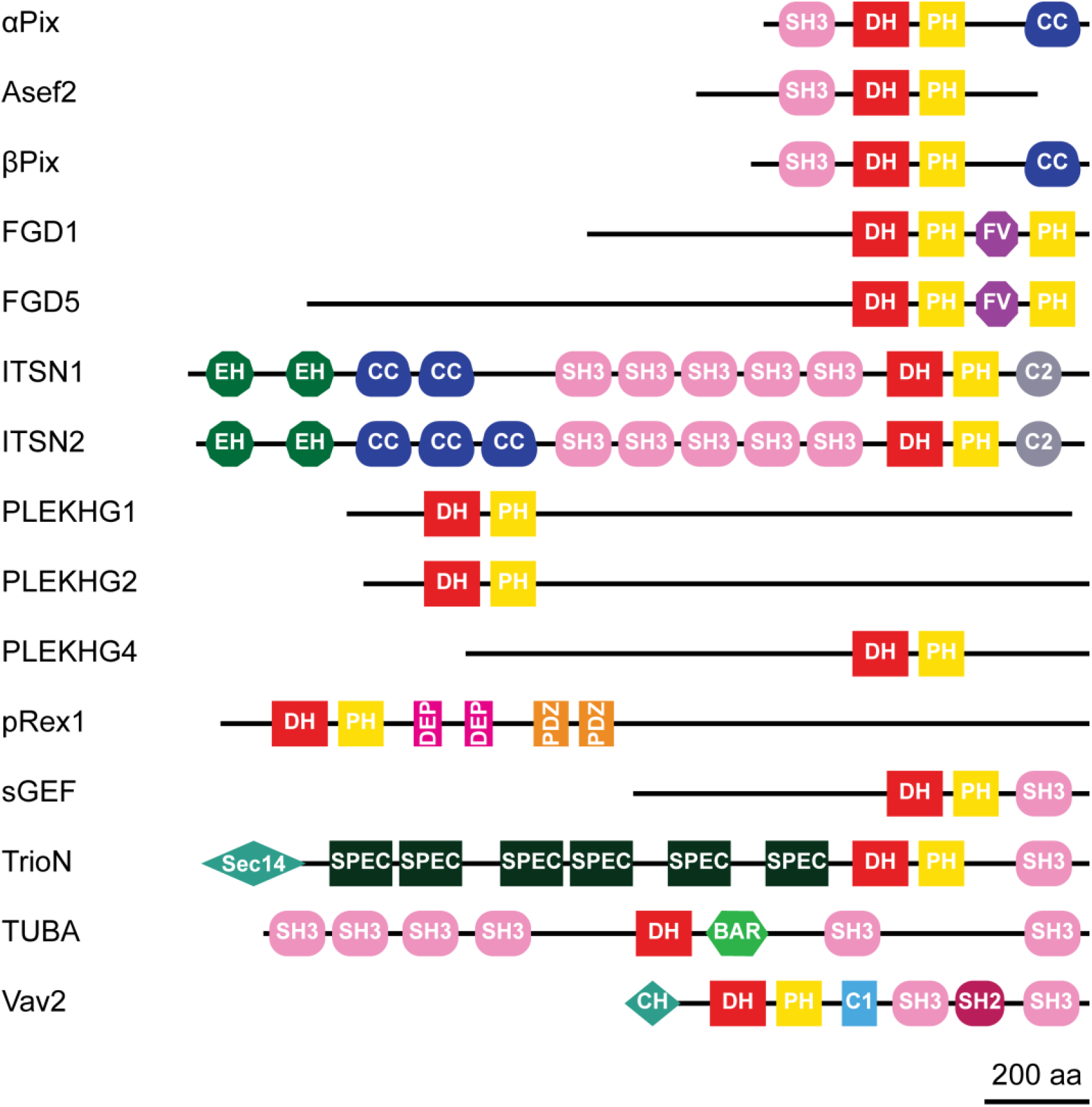
Protein domain structure of potential Cdc42 GEFs. Size of GEF structure relates to number of amino acids. Scale bar = 200 amino acids (aa). SH3 = Src homology 3; DH = Dbl homology; PH = Pleckstrin homology; CC = coiled coil; FV = FYVE domain; EH = Eps 15 homology; C2 = Protein kinase C conserved region 2; DEP = domain found in Dishevelled, EGL-10 and pleckstrin; PDZ = domain present in PSD-95, Dlg homologous region and ZO-1/2; Sec14 = domain in homologous of a S. serevisiae phosphatidylinositol transfer protein; SPEC = spectrin repeat; BAR = Bin-Amphiphysin-Rvs; CH = Calponin homology domain; C1 = Protein kinase C conserved region 1; and SH2 = Src homology 2.

### GEF-phenotypes in ECs

Most of the selected GEFs have not been previously studied in ECs. The initial characterization of their potential function entailed analysis of their intracellular localization. Unfortunately, due to the lack of proper antibodies, it is not possible to study localization of the endogenous proteins. Therefore, we generated GEF fusions with the cyan fluorescent protein mTurquoise2 (mTq2), to explore individual GEF localization.

Analysis of expression of mTq2-GEFs in confluent EC monolayers was combined with immunofluorescent staining for F-actin and the cell-cell contact protein Vascular Endothelial (VE)-cadherin (Figure 2A). This strategy induced notable phenotypes in ECs. A specific ‘protruding’ phenotype was observed for FGD1, PLEKHG2 and Vav2. In cells expressing these GEFs, the peripheral membrane protrudes beyond the VE-cadherin-positive cell-cell junction, a phenomenon that is also induced by the expression of constitutively active Cdc42 (Cdc42-G14V) (Figure 2B). FGD1 and Vav2 furthermore show a marked increase in cortical actin, while PLEKHG2 itself partly localized at F-actin fibers. In contrast to this protruding phenotype, expression of β-Pix resulted in a ‘contractile’ phenotype, marked by an increase in F-actin stress fibers and a jagged appearance of the VE-cadherin complex.

**Figure 2.**
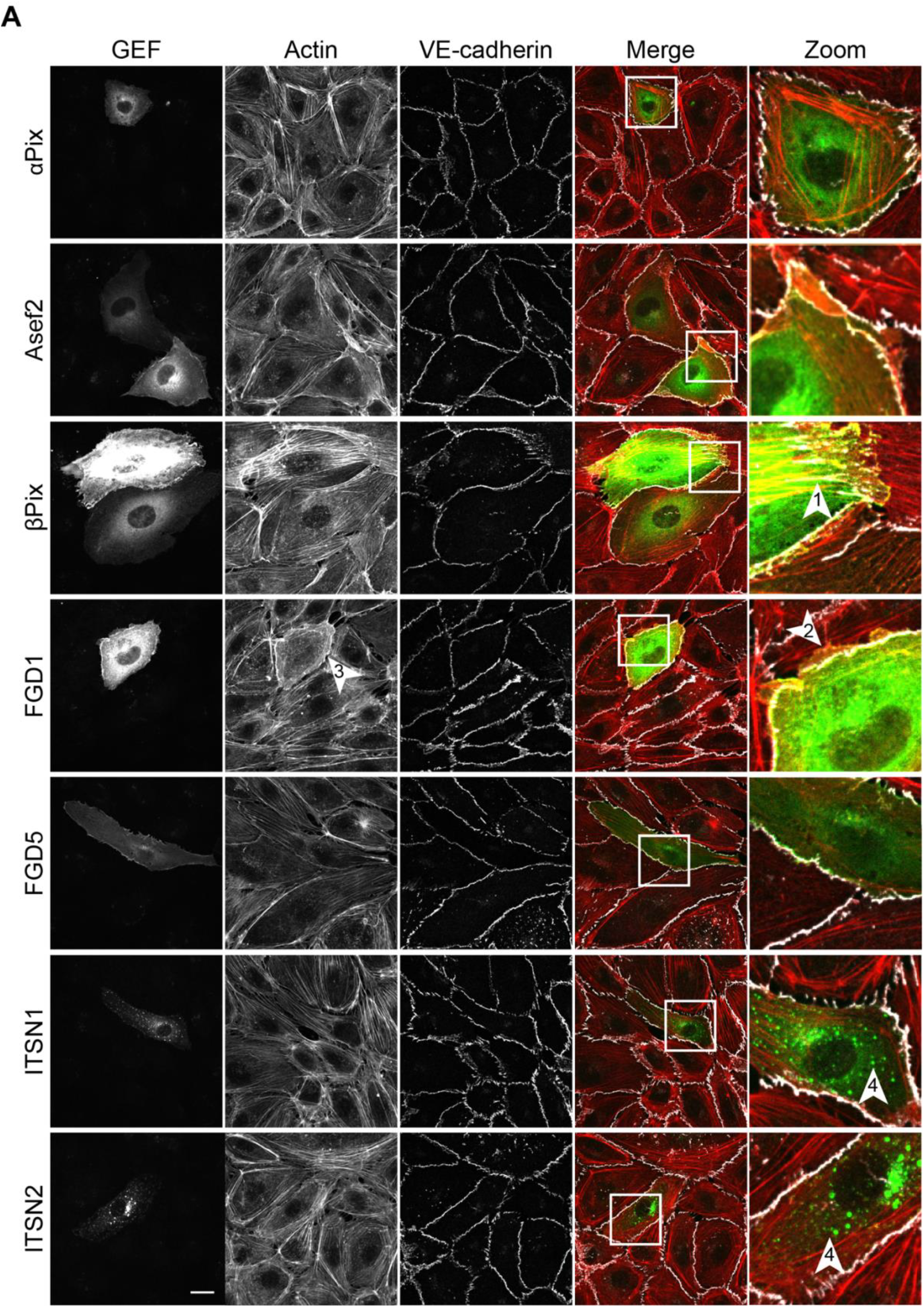
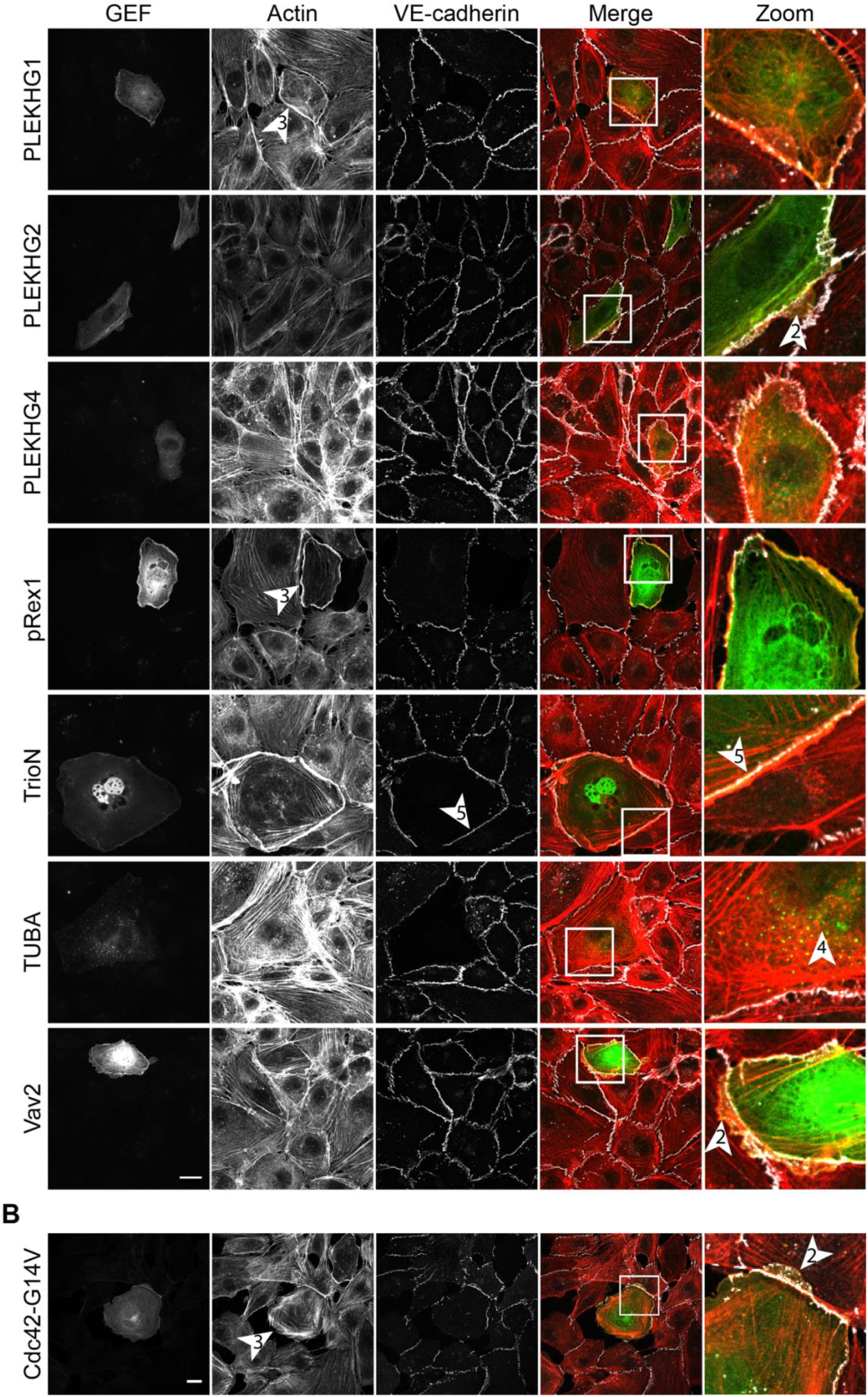
Ectopic expression of potential Cdc42 GEFs induces specific phenotypes in ECs. **A)** ECs were transiently transfected with mTq2-fused α-Pix, Asef2, β-Pix, FGD1, FGD5, ITSN1, ITSN2, PLEKHG1, PLEKHG2, PLEKHG4, pRex, TrioN, TUBA or Vav2. ECs were grown to a monolayer and stained for F-actin and VE-cadherin. Arrowheads highlight specific phenotypes: arrowhead #1 contractile phenotype, arrowhead #2 protruding phenotype, arrowhead #3 cortical actin, arrowhead #4 vesicle-like structures, arrowhead #5 lineair VE-cadherin. Except for the zoomed images, image acquisition and processing are equal between all conditions. **B)** ECs were transiently transfected with Cdc42-G14V, grown to a semi-confluent monolayer and stained for F-actin and VE-cadherin. Arrowhead #2 indicates protruding phenotype, Arrowhead #3 indicates cortical actin. Scale bars of **A** and **B** are 20 μm.

While most of the GEFs showed homogeneous localization throughout the cell, localization to vesicle-like structures was observed for ITSN1, ITSN2 and TUBA. Expression of PLEKHG1 and pRex1, induced a large fraction of cortical actin. Next to this induction of cortical actin, we observed dissociation from the cell matrix, combined with reduced levels of VE-cadherin at cell-cell junctions. TrioN positive cells, showed nuclear accumulation and a typical linear VE-cadherin phenotype. These linear VE-cadherin junctions have already been reported as a TrioN-specific phenomenon^23–25^. No specific phenotypes were observed for α-Pix, Asef2, FGD5 and PLEKHG4

Together, these data localize mTq2-tagged GEFs in human ECs and show that the GEFs induce specific and differential phenotypes, inferred from endogenous F-actin- and VE-cadherin labeling.

### A single-cell FRET-based approach to study GEF activation of Cdc42

The GEF localization approach revealed various GEF-specific phenotypes in ECs. Although some of these phenotypes hint towards activation of Cdc42, direct evidence is lacking. In order to study GEF-mediated Cdc42 activation directly, we applied a single cell FRET-biosensor imaging strategy, summarized in Figure 3. This strategy involves a FRET biosensor which records Cdc42 activation^26^. The sensor read-out uses a cyan fluorescent protein (CFP) and a yellow fluorescent protein (YFP) as a FRET pair that allows ratiometric image-based analysis, in which an increase in YFP/CFP ratio corresponds to an increase in Cdc42 activation.

**Figure 3.**
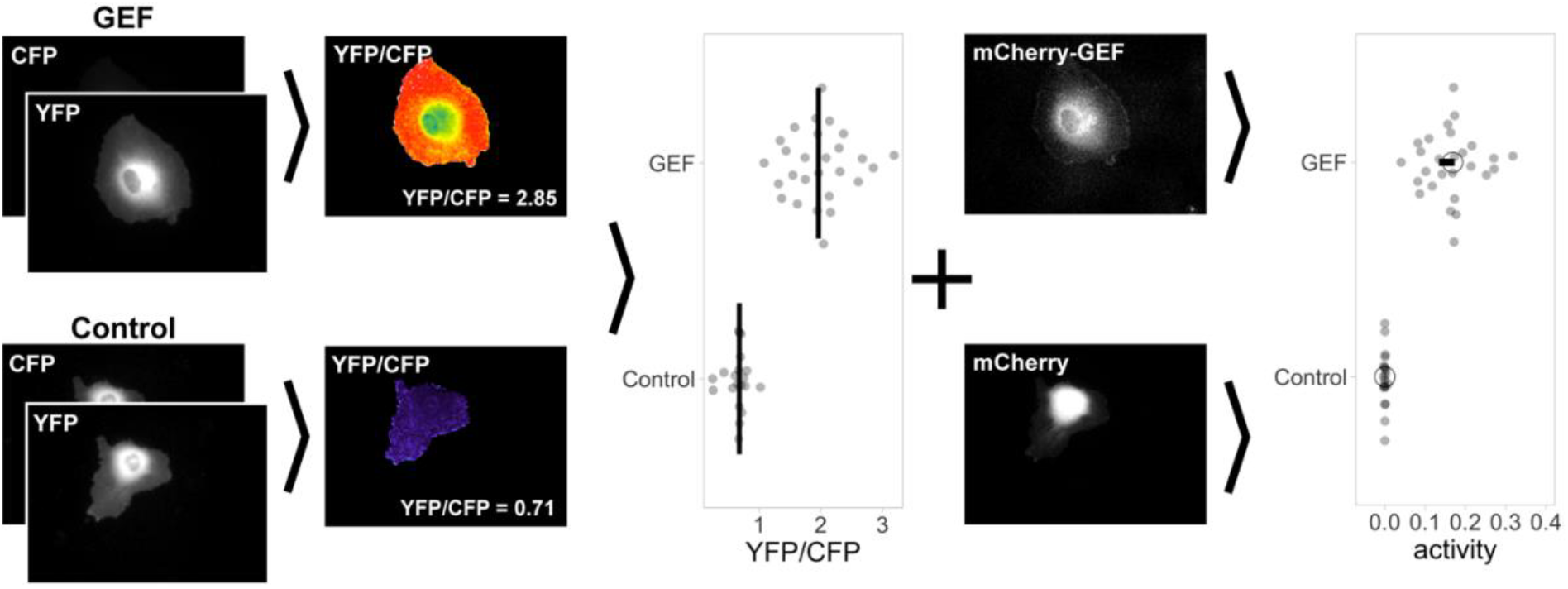
Workflow of GEF-mediated Cdc42 activation measurements in single ECs. Data points are part of an actual experiment included in this paper. ECs were transiently transfected with the YFP/CFP-based Cdc42 FRET sensor, and either C1-mCherry- or a mCherry-GEF. YFP and CFP images were acquired to generate the YFP/CFP ratio images and to quantify these values per cell. The median YFP/CFP value is depicted in the graphs as a vertical line. Subsequently, the mCherry intensity was used to convert the YFP/CFP ratio data into values of ‘relative activity’. The median activity is depicted as a circle and the 95% confidence interval determined by bootstrapping is indicated as a horizontal bar. For a detailed explanation see supplementary figure S1.

In our experimental setup, the Cdc42 sensor is co-expressed with either a soluble mCherry (as a control) or a mCherry-tagged GEF of interest. The fluorescence intensity of mCherry will reflect the concentration of the GEF. Cells that showed both sensor-as well as mCherry expression within the intensity range of 4-600, were selected. Next, image acquisition and processing resulted in a ratiometric YFP/CFP image, including a corresponding average YFP/CFP value. These average YFP/CFP values of multiple cells were plotted for both the control as well as for the GEF-expressing cells (Figure 3). In turn, YFP/CFP ratios and corresponding mCherry intensities were plotted in mCherry-YFP/CFP ratio graphs (supplementary figure S1). The rationale is that the Rho GTPase activity, assessed from the YFP/CFP ratio, is correlated with GEF activity, inferred from mCherry fluorescence. To quantify the relationship between activity and GEF concentration, the data points (each dot represents a single cell) were subjected to linear regression analysis via the Theil-Sen estimator method. The Theil-Sen estimator method calculates the median slope, which we take as a measure of relative activity of a GEF. For a quantitative comparison between conditions, we plot the median activity and the 95% confidence interval (Figure 3). Details of this Theil-Sen estimator analyses are summarized in Supplemental Figure S1.

In summary, the cell-based analysis of Rho GTPase activation with FRET based biosensors and tagged GEFs enables the determination and direct quantitative comparison of GEF activity in single living ECs.

### GEF expression induces specific Cdc42 and Rac1 activation patterns in ECs

The workflow described in the previous section was applied for all selected GEFs listed in Figure 1. First, the YFP over CFP ratios were determined for ectopically expressed GEFs. (Figure 4A). Next to Cdc42 activation also Rac1 activation was monitored, using a validated Rac1 FRET sensor^25^ (Figure 4B). The YFP/CFP ratios were subsequently converted into ‘relative activity’ values by taking GEF expression levels into account. The resulting activities are sorted according to their median value and plotted with a 95% confidence interval (Figure 4C and D). Based on arbitrarily set cut-off values we have defined three levels of activation of Cdc42 i) no Cdc42 activation; Asef2, β-Pix, α-Pix and FGD5 ii) intermediate activation; TUBA, Vav2, PLEKHG4, ITSN2, TrioN, sGEF and ITSN1, and iii) strong activation; PLEKHG2, FGD1, PLEKHG1 and pRex1. A similar categorization for Rac1 defines the GEFs with i) no Rac1 activation; FGD1, Asef2, β-Pix, α-Pix, FGD5 and ITSN1, ii) intermediate activation; PLEKHG1, PLEKHG4, sGEF, ITSN2, PLEKHG2 and TUBA, and iii) strong activation; Vav2, pRex1 and TrioN (Figure 4C, D).

**Figure 4.**
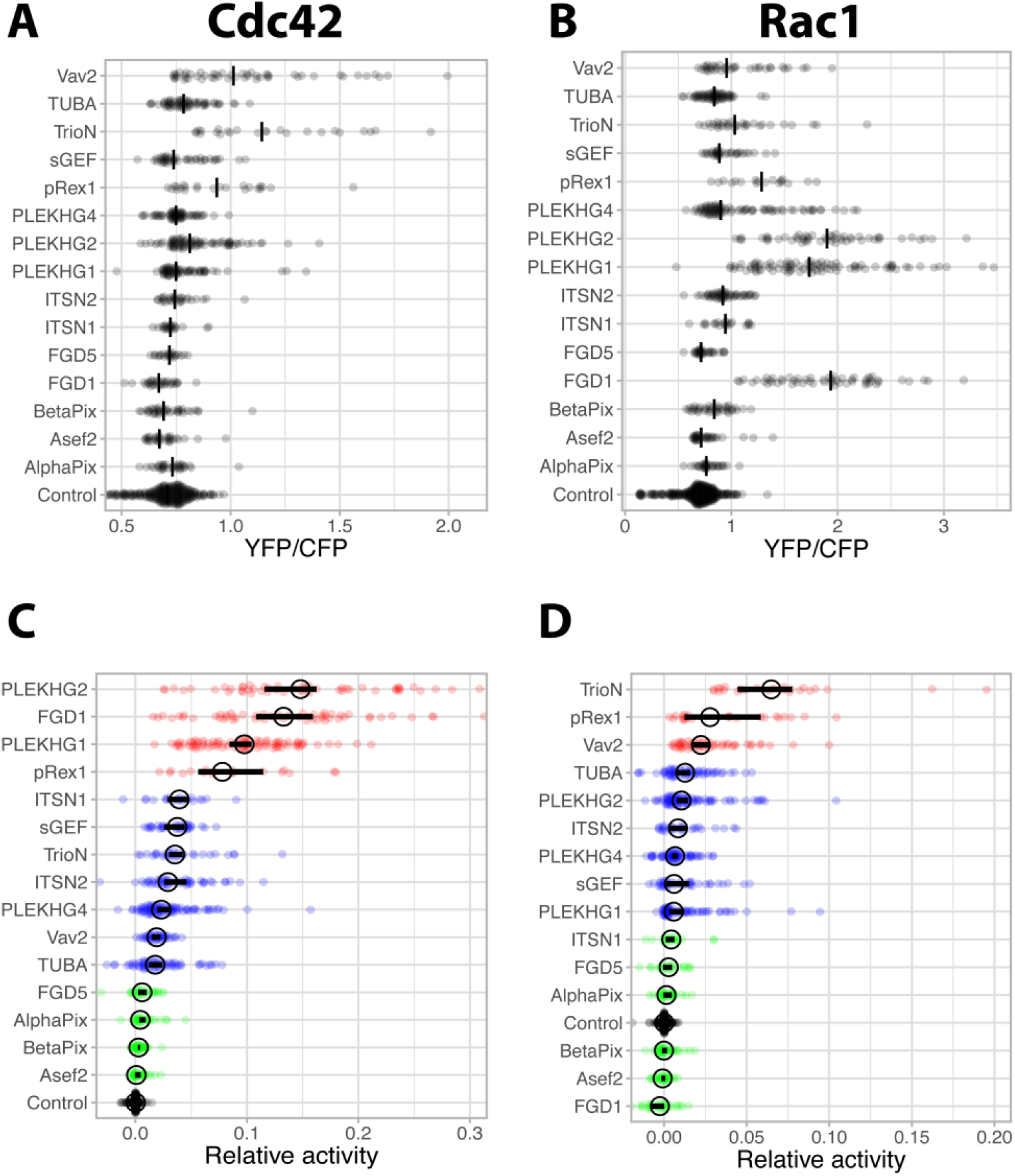
Ectopic expression of GEFs induce distinct Cdc42 and Rac1 activation patterns. **A)** Average YFP/CFP ratios of ECs that were transiently transfected with the Cdc42 FRET sensor and either C1-mCherry (Control, n=296) or mCherry-fused GEFs (α-Pix n=30, Asef2 n=33, β-Pix n=38, FGD1 n=55, FGD5 n=29, ITSN1 n=20, ITSN2 n=49, PLEKHG1 n=88, PLEKHG2 n=70, PLEKHG4 n=82, pRex1 n=18, sGEF n=33, TrioN n=31, TUBA n=52 and Vav2 n=36). **B)** Average YFP/CFP ratios of ECs that were transiently transfected with the Rac1 FRET sensor and either C1-mCherry (Control, n=271) or mCherry-fused GEFs (α-Pix n=33, Asef2 n=31, β-Pix n=38, FGD1 n=31, FGD5 n=22, ITSN1 n=21, ITSN2 n=28, PLEKHG1 n=64, PLEKHG2 n=81, PLEKHG4 n=61, pRex1 n=24, sGEF n=41, TrioN n=24, TUBA n=61 and Vav2 n=50). **C&D)** The relative activity calculated from the YFP/CFP ratios and the mCherry intensities quantified from single cells. The median and 95% confidence interval are indicated by a circle and horizontal bar respectively. The green, blue and red colors define no activation, intermediate activation and strong activation respectively. Online, interactive plots are available for <u>panel C</u> and <u>panel D</u>.

Overall, these data show distinct GEF-induced Cdc42 and Rac1 activation profiles in live ECs. Different GEFs can be divided into low-, medium-, and high activators, with differential preference for Cdc42 or Rac1 GTPases.

### TIAM1 exclusively activates Rac1, and not Cdc42

As shown in Figure 4, pRex1, TrioN and Vav2 all, to some extent, activate Cdc42. Previous literature, however, mainly describes these GEFs as Rac1 activators. To test the specificity and discriminative power of our assay, we used the GEF TIAM1 as an additional negative control. Based on vitro data ^13^, TIAM1 is highly selective, showing high GEF activity towards Rac1, but not towards Cdc42. Applying the same strategy as described in Figure 3, the catalytic domain of TIAM1 was used to study the effect on Cdc42 or Rac1. Ectopic expression of mCherry-TIAM1, combined with the Cdc42 FRET sensor, did not show Cdc42 activation based on the YFP/CFP ratio (Figure 5A) and also its relative activity was near the control value (Figure 5C). In contrast to Cdc42, strong effects were observed when the Rac1 sensor was used in combination with TIAM1. Both the YFP/CFP ratios as well as the relative activity were strongly increased relative to the control (Figure 5B&D). These data suggest that the Cdc42 biosensor is not sensitive for Rac GEFs and that induction of Rac1 activity does not (indirectly) result in Cdc42 activation in endothelial cells.

**Figure 5.**
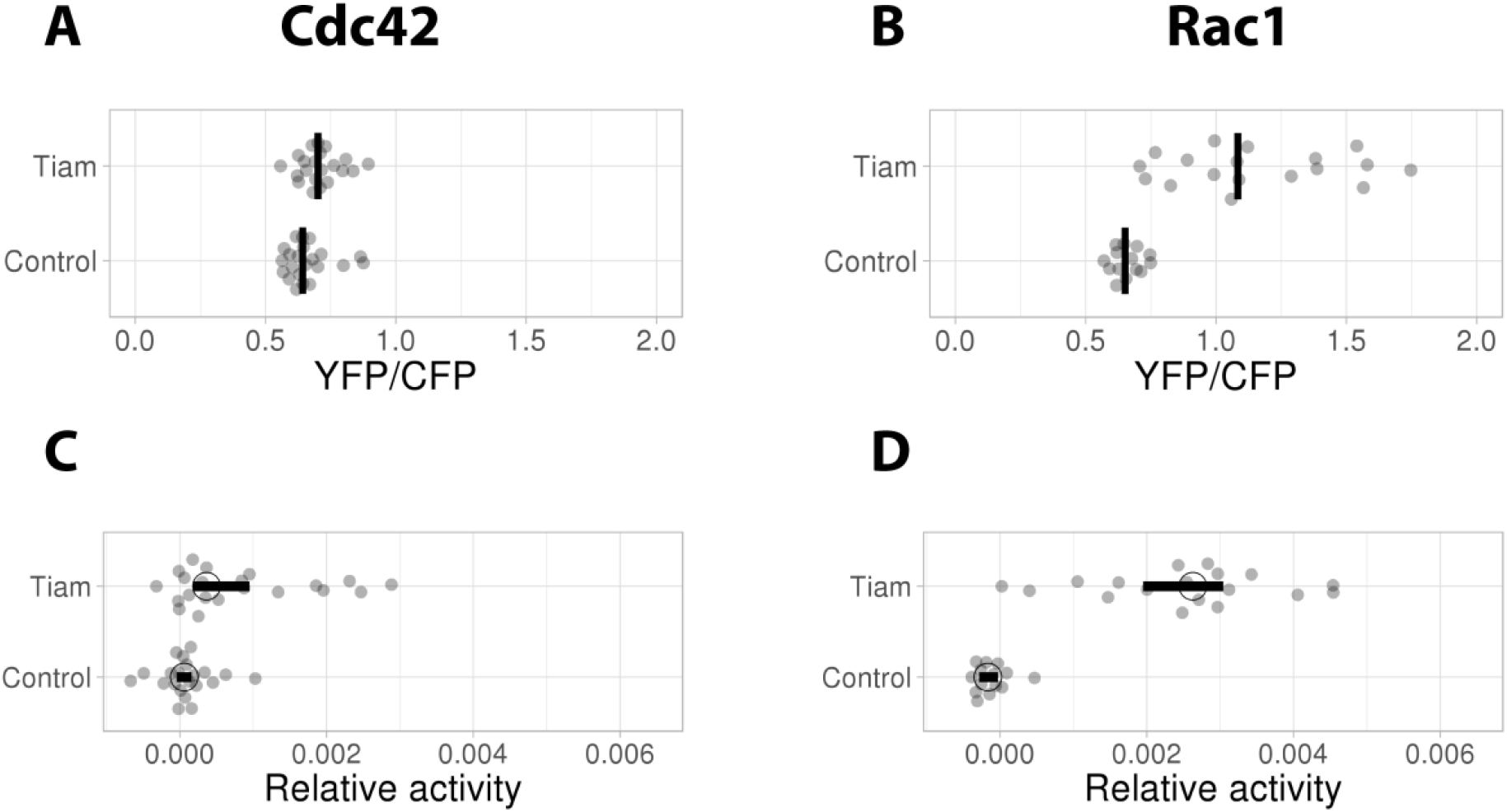
Ectopic expression of TIAM exclusively activates Rac1. **A)** YFP/CFP ratios of ECs that were transiently transfected with the Cdc42 FRET sensor and either C1-mCherry (Control, n=22) or mCherry-TIAM (n=21). **B)** YFP/CFP ratios of ECs that were transiently transfected with the Rac1 FRET sensor and either C1-mCherry (Control, n=14) or mCherry-TIAM (n=18). **C)** The relative activity observed with the Cdc42 sensor based on the YFP/CFP ratios shown in panel A and mCherry intensity. **D)** The relative activity observed with the Rac1 sesnor based on the YFP/CFP ratios shown in panel B and mCherry intensity. In C and D the median and 95% confidence interval are indicated with a circle and bar respectively.

Collectively, our data demonstrate that the Cdc42 and Rac1 biosensors report with high selectivity on GEF activity in a cellular context.

### Catalytic GEF domains induce differential Cdc42 activation patterns

As can be inferred from the protein domain structures in Figure 1, the GEFs of interest belong to the Dbl family (containing a DH and PH domain). The activity that we have measured for the FL GEFs (Figure 4) is a basal activity determined by the combined role of all the domains that are present. Previous studies have demonstrated that the DH domain is responsible for the catalytic activity towards their preferred targets^4,10,13^. The PH domain that is adjacent to the DH domain in the majority of GEFs has a less clearly defined role. Finally, domains beyond the DHPH tandem can have diverse regulatory roles, including localization, autoinhibition, protein-protein or protein-lipid interactions. To study the role of the catalytic unit in more detail, we selected a number of GEFs to examine the activity of the DH domain and the role of the PH domain.

In total eight GEFs from all three levels of Cdc42 activation (weak, intermediate, strong) were selected, including FGD1, FGD5, ITSN1, ITSN2, PLEKHG1, PLEKHG2, PLEKHG4 and pRex1. Corresponding GEF DHPH domain alignments (acquired from ClustalW) highlight the amino acid sequence homology between these proteins (Figure 6). We generated mCherry-fused DH, and DHPH constructs. Since it has been shown that recruitment of a GEF to the plasma membrane is sufficient to increase Rho GTPase activity, we also made variants with a plasma membrane targeting signal from Lck.

**Figure 6.**
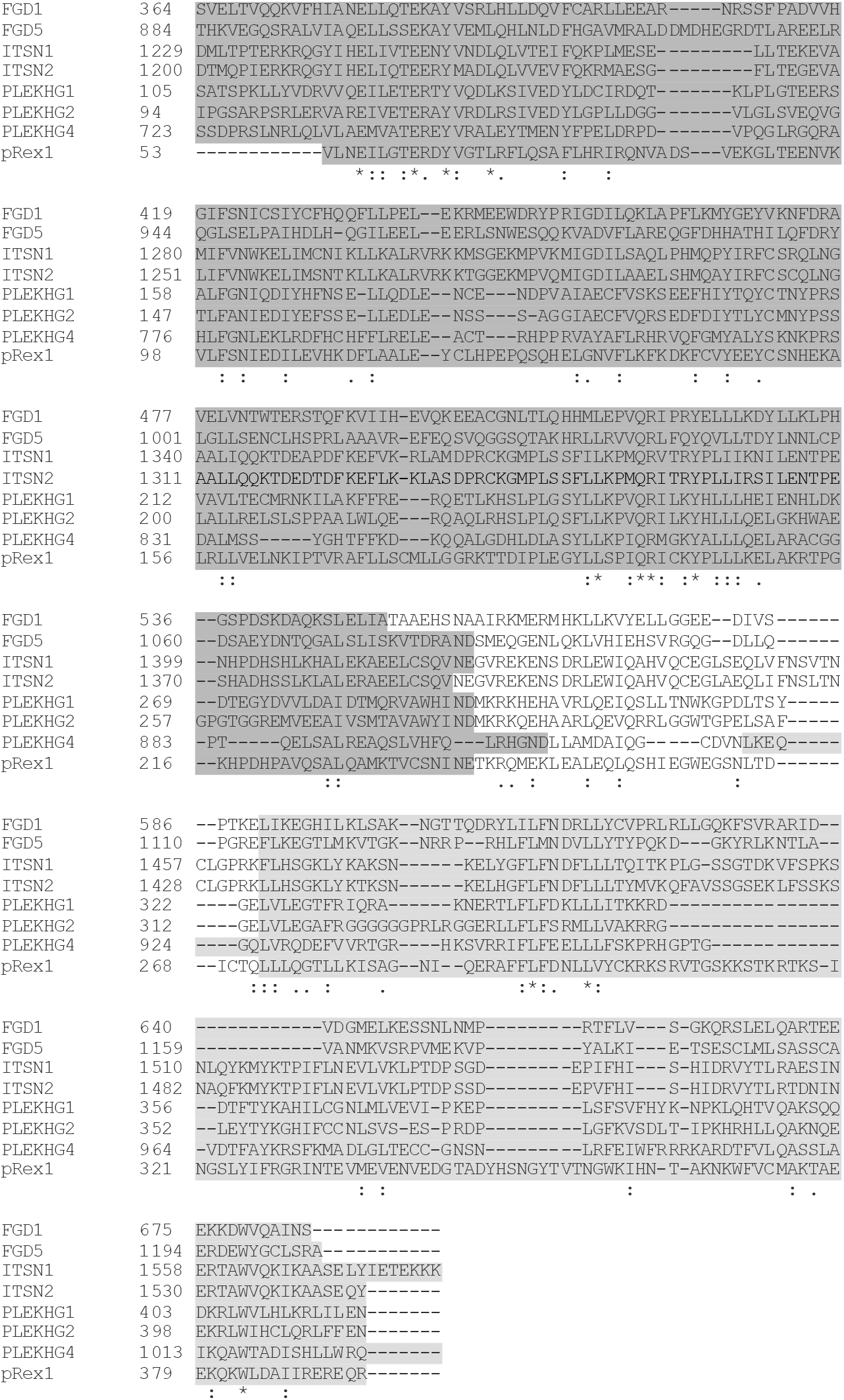
Alignment of GEF DHPH domains. Alignment is performed on FGD1, FGD5, ITSN1, ITSN2, PLEKHG1, PLEKHG2, PLEKHG4 and pRex1. Numbers indicate corresponding amino acid position of the FL protein. Dark gray represents the DH domain, light gray represents the PH domain. **\*** represents single, fully conserved residue, : represents conserved residues between groups of strongly similar properties,. represents conserved residues between groups of weakly similar properties.

We tested to what level the different DH/DHPH constructs activated Cdc42, relative to the corresponding full length (FL) GEFs. Distinct activation patterns were observed for the GEFs of interest (Figure 7A-H and Supplemental Figure S2). PLEKHG2 and pRex1 showed pronounced Cdc42 activation by the FL protein, while the DH/DHPH constructs hardly showed Cdc42 activation (Figure 7F, H and Supplemental Figure S2F, H). In contrast to the average YFP/CFP ratio plots, a pronounced effect was observed for the relative activities of three of the FGD1 truncation constructs; DH, DHPH and Lck-DHPH, respectively (Figure 7A and Supplemental Figure S2A). We think that the relative activity measure is a better indicator of activating potential of a GEF, since it takes the expression level of the GEF into account.

**Figure 7.**
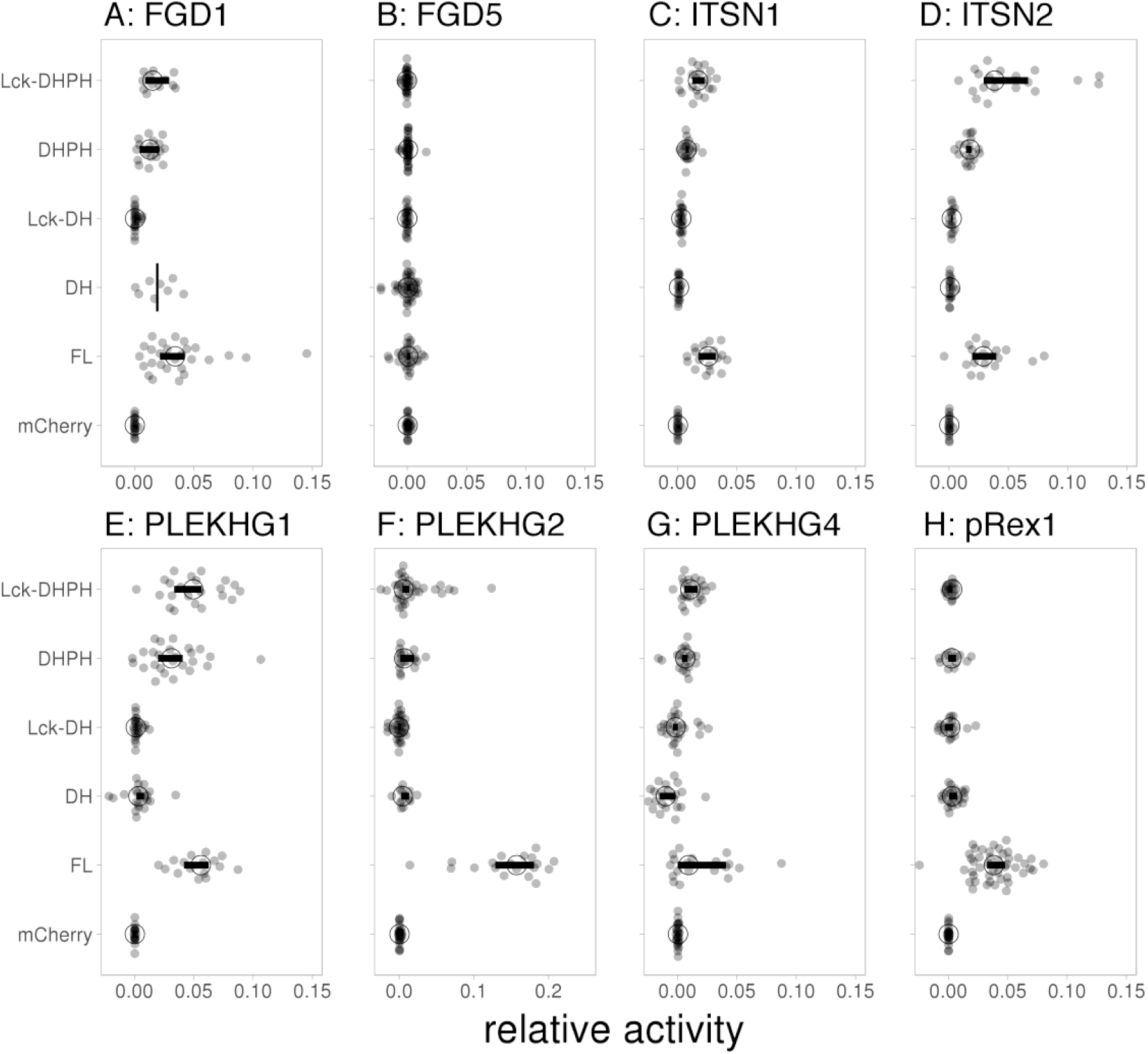
Catalytic GEF domains induce distinct Cdc42 activation profiles. For each of the indicated GEFs the relative activity of membrane targeted (Lck) Lck-DHPH, soluble DHPH, Lck-DH, soluble DH or full-length (FL) on the Cdc42 biosensor was quantified relative to the control (mCherry). The median activity and 95% confidence intervals are indicated with a circle and a horizontal bar respectively (except for the ‘DH’ condition of FGD1, where only the median is indicated due to low sample size). The corresponding YFP/CFP data are represented in Supplemental Figure S2.

A difference in YFP/CFP ratio and activity value was not observed for PLEKHG2 and pRex1 (Figure 7F, H and Supplemental Figure S2F, H). For the weak activator FGD5, neither the average YFP/CFP ratio, nor the activity values were affected in any of the conditions (Figure 7B and Supplemental Figure S2B). Relative to the FL and the DH constructs, the DHPH constructs induced high YFP/CFP ratio increases for ITSN1, ITNS2, PLEKHG1 (Suppemental Figure S2C, D, E). A different pattern was observed in the corresponding relative activities. Here the effect of the DHPH truncation constructs was less pronounced relative to the FL constructs (Figure 7C, D, E and Supplemental Figure S2C, D, E). Finally, the catalytic PLEKHG4 constructs induced moderate increases in Cdc42 activation as compared to the FL PLEKHG4 (Figure 7G and Supplemental Figure S2G)

In summary, For FGD1, ITSN1, ITSN2 and PLEKHG1 different patterns were observed when comparing average FRET ratios and corresponding relative activities. Together, our data show that catalytic domains of different GEFs, show unanticipated and distinct Cdc42 activation profiles.

## Discussion

Over the past years more than 80 GEFs have been identified and they have been shown to be involved in a variety of functions and signaling networks. This study focusses on a selection of Dbl-family RhoGEFs, that may signal towards Cdc42 in the endothelium, including α-Pix, Asef2, β-Pix, FGD1, FGD5, ITSN1, ITSN2, PLEKHG1, PLEKHG2, PLEKHG4, pRex1, sGEF, TrioN, TUBA and Vav2. Fluorescent-labeled versions of these GEFs provide new insights regarding the localization of these proteins in primary human ECs. We furthermore implement a robust FRET-based and live-cell assay, to measure and quantitatively compare GEF-mediated activation of Cdc42. This demonstrates differential activation patterns, induced by FL GEFs or their isolated DH/DHPH domains.

So far, most of these selected GEFs have not been studied in the endothelium. We visualize these proteins in ECs by generating mTq2-labeled GEF constructs and reveal diverse localizations and different GEF-induced phenotypes. Although these localization studies are descriptive, specific phenotypes (e.g. FGD1-induced membrane protrusions) already hint towards the activation of Cdc42.

Integrating GEF overexpression with our single-cell FRET strategy, allowed us to generate large datasets of GEF-induced activation of Cdc42 or its close homologue Rac1. To relate the observed activities to GEF expression levels, we performed Theil-Sen estimation analyses, generating slope values for each individual data point. The outcome of the analysis, the relative activity, reflects the intrinsic, basal activity of the GEF. Based on this strategy, we observe differential activation profiles for Cdc42 and Rac1, underscoring the specificity of this assay. Of note, the amplitude differences in YFP/CFP ratio between Rac1 and Cdc42 biosensors are most likely the result of a higher dynamic range of the Cdc42-FRET sensor. This technical limitation still allows us to compare trends in Cdc42 and Rac1 activation while comparisons of actual YFP/CFP ratios and absolute activity values are not straightforward.

The strongest Cdc42 activators comprise FGD1, PLEKHG1 and PLEKHG2. The lack of FGD1-induced Rac1 activation underscores the specificity of FGD1-induced Cdc42 activation. This is well in line with previous studies that characterized FGD1 as a Cdc42 GEF involved in faciogenital dysplasia, a human disease that affects skeletogenesis^27,28^. We observe most prominent Rac1 activation for TrioN, pRex1, Vav2 and TIAM1, in line with previous studies which defined these as Rac1 GEFs ^13^. Mainly for pRex1, and to some extent for TrioN and Vav2, we could also detect elevated Cdc42 activation. Strikingly, the activity of pRex1 towards Cdc42 has been observed using purified components^13^, but not in cellular context^29^. Whether It is unlikely that the effect on Cdc42 is indirect, since overexpression of the Rac GEF TIAM did not result in Cdc42 activation the effects that we observe are indirect, i.e. through Rac1, or direct requires further molecular analysis. While FGD5, ITSN1 and TUBA have already been linked to Cdc42 activation^30–32^, our approach only detects moderate signaling towards Cdc42. However, it should be noted that our approach measures basal activity of the GEFs. The activity may be increased by signaling or affected by the binding of accessory/scaffolding proteins. A limitation of our assay is that we may miss the effects of endogenous accessory proteins due to over over-expression of the GEFs. Additionally, In ECs, we observe that ITSN1 and TUBA primarily localize at vesicles. But, it is unexplored whether Cdc42 can get activated at these localized structures.

GEF activities of the full-length proteins can be influenced by auto-inhibitory domains^33^, protein-protein interactions and by phosphorylation of specific residues^34^. Although FL GEF studies may appear physiologically the most relevant, the use of isolated GEF domains limits complexity. A selection of our catalytic constructs induces elevated Cdc42 activation, most clearly for the membrane-linked, DHPH constructs of ITSN1, ITSN2 and PLEKHG1 and to some extent for FGD1 and PLEKHG4. This not only demonstrates the functionality of the respective DHPH domains, but also proposes these GEFs as interesting candidates in recruitment- and/or optogenetic systems, to locally induce Cdc42 activation at the plasma membrane. We^35^, and others^36,37^, have already demonstrated the value of these approaches in biological systems. Interestingly, we observe distinct trends in activation when comparing YFP/CFP ratio- and corresponding activity analyses. Compared to DHPH domains, FL GEFs exhibit higher activation potencies measured by ‘relative activity’ as compared to YFP/CFP ratio analysis. This implies that the power of catalytic GEF-mediated Cdc42 activation largely depends on the expression level of respective DHPH domains.

Previously, we observed with a similar strategy that the PH domain of p63RhoGEF has an autoinhibitory role and that the isolated DH domain has high activity towards RhoA^35^. However, we did not find a similar role for the PH domain in any of the GEFs that we analyzed. Strikingly, the isolated DH domains (with the exception of the DH domain of FGD1) did not show any activity in a cellular context. It appears that the PH domains of the Cdc42 GEFs that we selected have a stimulatory rather than inhibitory role.

In contrast to the previous-mentioned GEFs, effects of DHPH domains of FGD5, PLEKHG2 and pRex1 are either small or absent. For these GEFs, additional domains could be required for full activity; this will require additional domain- and activity analyses.

In conclusion, this study provides new insights regarding GEF-mediated Cdc42 activation in live, primary human ECs. Our FRET-based GEF screening method identifies FL PLEKHG2, FGD1, PLEKHG1 and pRex1 as prominent Cdc42 activators. Additionally, catalytic domains of ITSN1, ITSN2 and PLEKHG1 behave as potential activators for local-induced Cdc42 activation at the plasma membrane. Overall, these findings uncover novel potential Cdc42-related (signaling) studies in the endothelium.

## Methods

### Plasmids

#### FL GEFs

For Asef2 (kind gift from D. Webb) and pRex1 (kind gift from H.C. Welch), mTq2 and mCherry were amplified by PCR. PCR products (inserts) and vectors were cut with restriction enzymes. Next, digested products were ligated to generate mTq2- and mCherry GEF fusions.

For β-Pix and TrioN (kind gifts from J. van Buul), vectors and GEF constructs were cut with restriction enzymes Next, digested products were ligated to generate mTq2- and mCherry GEF fusions.

For α-Pix (kind gift from J.L. Zugaza), FGD1 (kind gift from M. Hayakawa), FGD5 (kind gift from W.J. Pannekoek), ITSN1 (obtained from addgene, plasmid #47395), ITSN2 (kind gift from I.G. Macara), PLEKHG1 (kind gift from K. Mizuno), PLEKHG2 (kind gift from H. Ueda), PLEKHG4 (kind gift from D. Manor), sGEF (kind gift from J. van Buul), TUBA (kind gift from L.J.M. Bruurs) and Vav2 (obtained from Addgene, plasmid #14554), GEFs were amplified by PCR. PCR products (inserts) and vectors were cut with restriction enzymes. Next, digested products were ligated to generate mTq2- and mCherry GEF fusions.

Corresponding PCR primers, restriction enzymes, and cloning products are listed in Table 1.

**Table 1.**
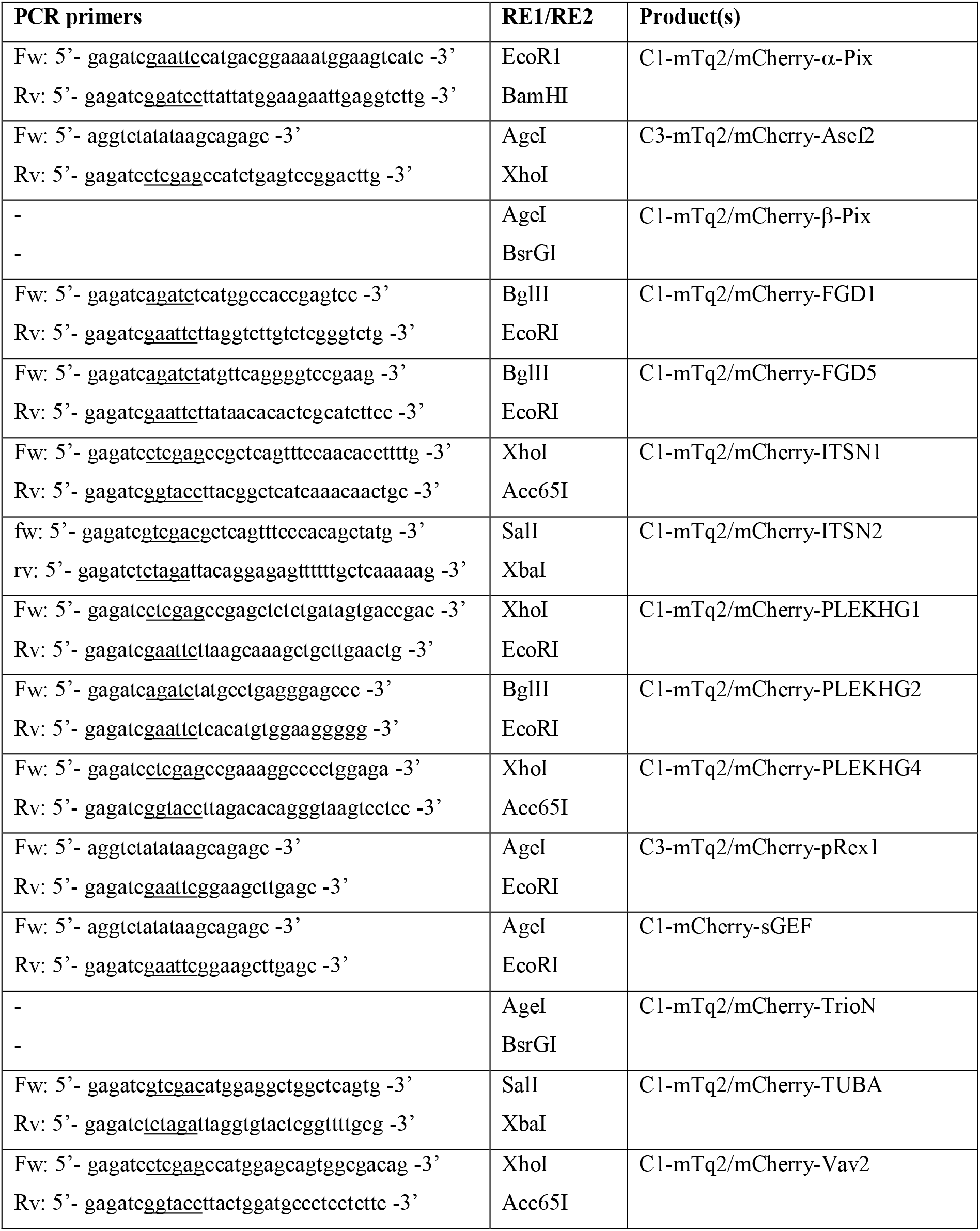
Generation of FL mTq2/mCherry-GEF constructs. RE = restriction enzyme, Fw = Forward primer, Rv = Reverse primer. Restriction sites are underlined in the primer sequences. Restriction sites for Asef2, pRex1 and sGEF are not present in the Fw primer.

#### DH(PH) truncation constructs

DH and DHPH domains of FGD1, FGD5, ITSN1, ITSN2, PLEKHG1, PLEKHG2, PLEKHG4 and pRex1 were amplified by PCR, using the same template constructs as for the cloning of the FL constructs. Next, PCR products (inserts) and vectors were cut with restriction enzymes. Finally, inserts were ligated into vectors to generate for each GEF mCherry-DH, Lck-mCherry-DH, mCherry-DHPH and Lck-mCherry-DHPH constructs. An overview of PCR primers and restriction enzymes are listed in Table 2.

**Table 2.**
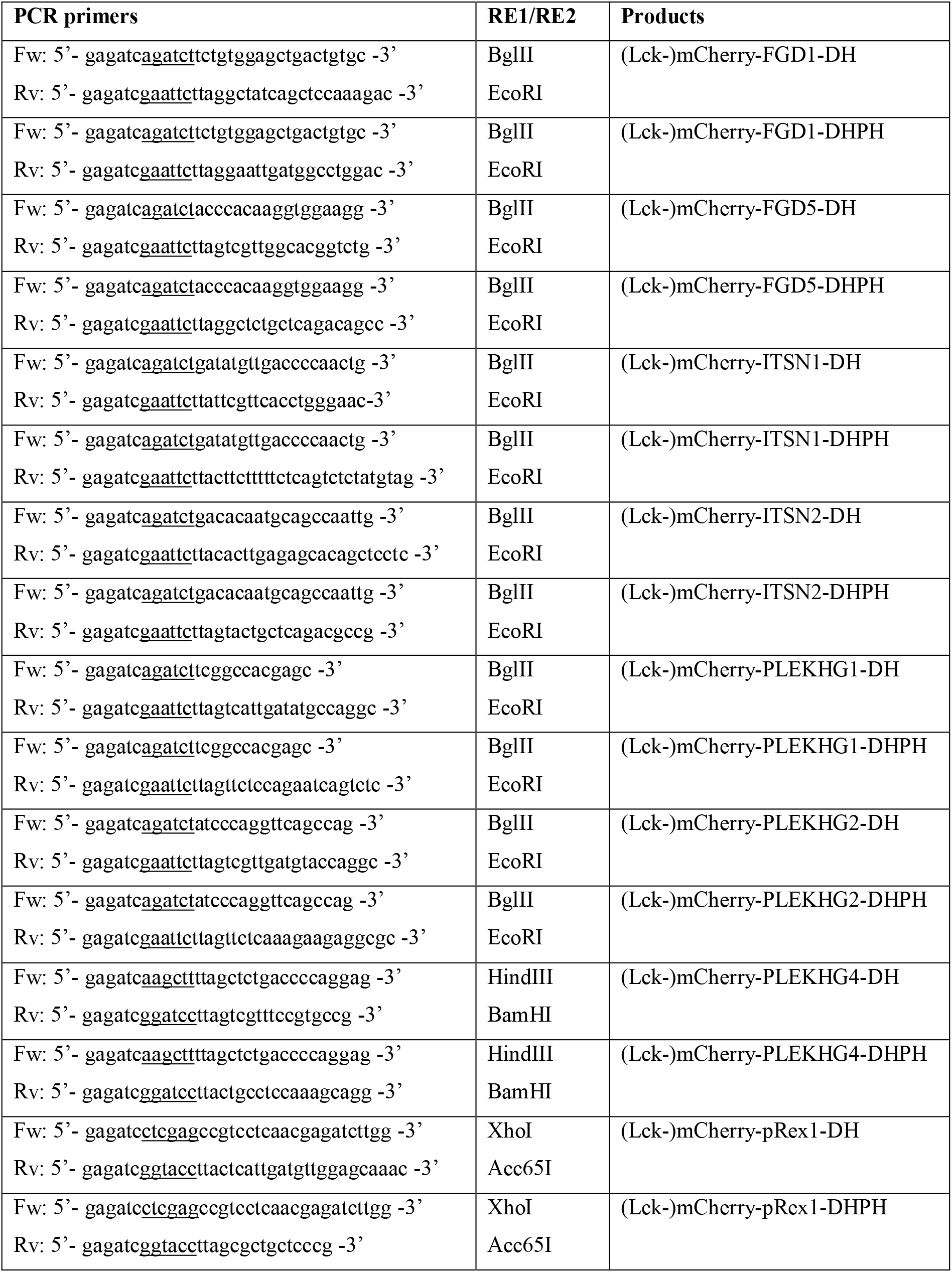
Generation of (Lck-)mCherry-DH(PH) GEF constructs. RE = restriction enzyme, Fw = Forward primer, Rv = Reverse primer. Restriction sites are underlined in the primer sequences.

#### Others

Cdc42-G14 (cDNA.org) was amplified by polymerase chain reaction (PCR), using forward primer (BsrGI site in uppercase): gcTGTACAagtccatgcagacaattaagtgtgt and reverse primer 3’- pcdna: gtcgaggctgatcagcgg. PCR product was digested with BsrGI and XbaI and inserted in a clontech style backbone. The Rac1 FRET sensor^25^ and Cdc42 FRET sensor^26,38^ were as published in the cited references.

### Immunofluorescence

Actin-stain 555 Phalloidin was from Cytoskeleton, monoclonal antibody (mAb) Mouse anti-VE-cadherin/CD144 AF647 was purchased from BD Pharmingen

### HUVEC culture and transfection

Primary HUVECs, purchased from Lonza (Verviers, Belgium), were seeded on fibronectin (FN)-coated culture dishes and grown in EGM2 medium (supplemented with singlequots (Lonza)). HUVECs (at passage #4 or #5) were transfected with 2μg plasmid DNA, using a Neon transfection system (MPK5000, Invitrogen) and a corresponding Neon transfection kit (Invitrogen) that generates a single pulse at 1300 V for 30 ms. After microporation, HUVECs were seeded on FN-coated glass coverslips.

### Confocal imaging

Transfected HUVECs were grown to a monolayer, washed in PBS (1mM CaCl_2_ and 0.5 mM MgCl_2_) and fixed in a PBS solution (1mM CaCl_2_ and 0.5 mM MgCl_2_) with 4% formaldehyde. After fixation, HUVECs were permeabilized for 5 min in PBS containing 0.5% Triton X-100 and blocked for 20 min in PBS containing 0.5% Bovine serum albumin (BSA). Finally, HUVECs were incubated for 1 hour with directly-labeled antibodies, dissolved in 0.5% PBS-BSA. Confocal images were obtained on a Nikon A1 confocal microscope, equipped with a 60x oil immersion objective (NA 1.40, Plan Apochromat VC) and Nikon NIS elements software.

### Live HUVEC FRET measurements

Glass coverslips with transfected HUVECs were mounted in Metal Attofluor cell chambers at least 16 hours after transfection. Live-cell FRET acquisitions were performed on a widefield Zeiss Axiovert 200 M microscope, equipped with an 40x oil-immersion objective (NA 1.30), Metamorph 6.1 software, a xenon arc lamp with mono-chromator (Cairn Research, Faversham, Kent, UK) and a cooled charged-coupled device camera (Coolsnap HQ, Roper Scientific, Tucson, AZ, USA). Cells were excited by using 420 nm light (slit width 30 nm) and a 455 DCLP (dichroic long-pass) mirror. Via a rotating filter wheel, CFP emission was directed to a 470/30 nm emission filter and YFP emission to a 535/30 nm emission filter. mCherry was excited with 570 nm light (slit width 10), and via a 585 DCXR mirror mCherry emission was directed to a 620/60 emission filter.

### Data analysis

All image acquisitions were background corrected and YFP acquisitions were bleed-through corrected (55% leakage of the CFP into the YFP channel). FRET images in Figure 3 were obtained by ImageJ, according to refs ^25,39^.

YFP/CFP ratio analysis was performed in Matlab (Matlab, The MathWorks, Inc., Natick, Massachusetts, United States). The Matlab script was first described in ref ^35^. Theil-Sen estimation analysis was performed according to Supplemental Figure S1. Of note, median slopes were determined and shown in the graphs, since median values are less prone to outliers. The graphs were made with PlotsOfData, showing the data as jittered dots, the median YFP/CFP values as a line and the median ‘activity’ values as a circle with 95% confidence interval indicated by a bar. The 95% confidence interval for the median was obtained by bootstrapping (1000 samples).

## Data availability

The data generated during this study is available at Zenodo.org: https://zenodo.org/record/2548920

## Acknowledgments

We thank D. Webb (Vanderbilt University), H.C. Welch (Babraham Institute), J. van Buul (Sanquin Research), J.L. Zugaza (University of the Basque Country), M. Hayakawa (Tokyo University of Pharmacy and Life Sciences), W.J. Pannekoek (University Medical center Utrecht), I.G. Macara (Vanderbilt University), K. Mizuno (Tohoku University), H. Ueda (Gifu university), D. Manor Case Western Reserve University, and L.J. Bruurs (University Medical center Utrecht) for providing us with the FL GEF constructs.

## Supplemental material

**Supplemental Figure S1.**
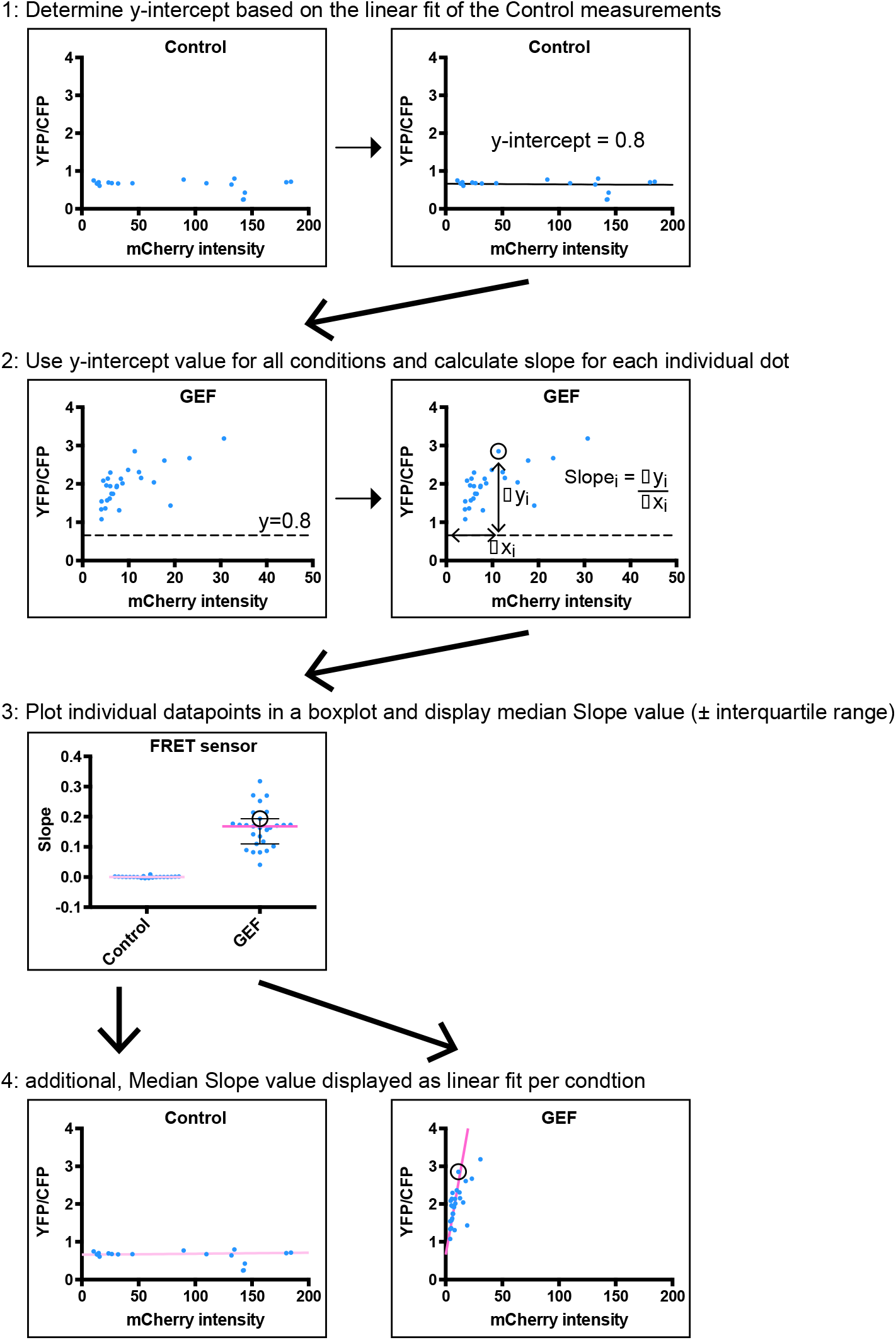
Workflow of Theil-Sen estimation method to calculate individual slopes based on mCherry-intensity -YFP/CFP graphs. Data points are part of an actual experiment included in this paper. The slope values that are calculated are used as a measure of ‘relative activity’ since these take the concentration of the GEF (through mCherry fluorescence intensity) into account

**Supplemental Figure S2.**
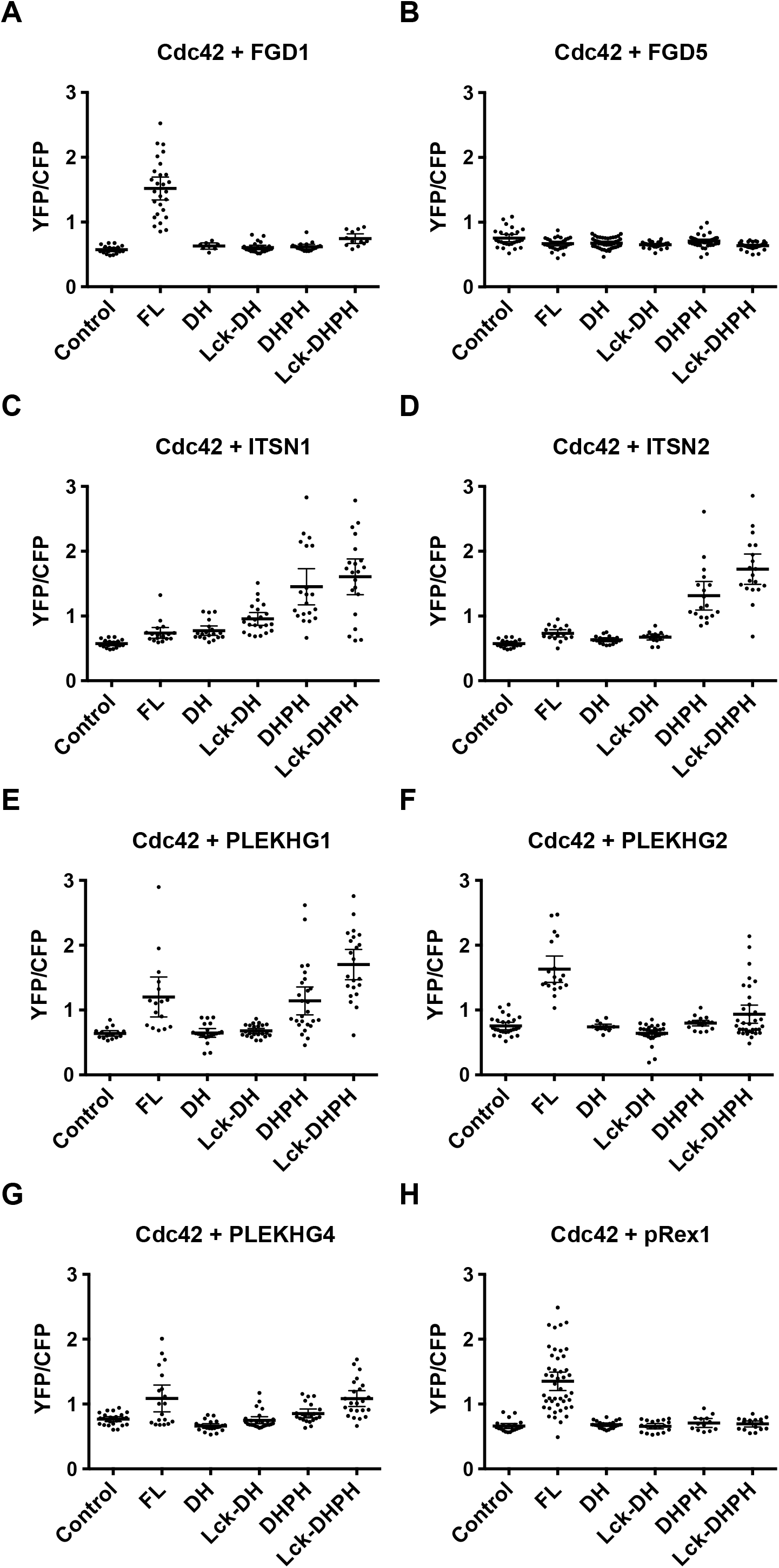
Catalytic GEF domains induce distinct activation profiles, based on median Slope values. Average YFP/CFP ratios (±95% CI) of ECs that were transfecte with either **A)** C1-mCherry (Control, n=18), mCherry-FGD1 (FL, n=27), mCherry-DH (DH, n=8), Lck-mCherry-DH (Lck-DH, n=23), mCherry-DHPH (DHPH, n=17), or Lck-mCherry-DHPH (Lck-DHPH, n=12). For **B)** with either C1-mCherry (Control, n=26), mCherry-FGD5 (FL, n=30), mCherry-DH (DH, n=40), Lck-mCherry-DH (Lck-DH, n=25), mCherry-DHPH (DHPH, n=40), or Lck-mCherry-DHPH (Lck-DHPH, n=33). For **C)** with either C1-mCherry (Control, n=18), mCherry-ITSN1 (FL, n=18), mCherry-DH (DH, n=19), Lck-mCherry-DH (Lck-DH, n=22), mCherry-DHPH (DHPH, n=20), or Lck-mCherry-DHPH (Lck-DHPH, n=21). For **D)** with either C1-mCherry (Control, n=18), mCherry-ITSN2 (FL, n=17), mCherry-DH (DH, n=18), Lck-mCherry-DH (Lck-DH, n=16), mCherry-DHPH (DHPH, n=18), or Lck-mCherry-DHPH (Lck-DHPH, n=19). For **E)** with either C1-mCherry (Control, n=17), mCherry-PLEKHG1 (FL, n=16), mCherry-DH (DH, n=21), Lck-mCherry-DH (Lck-DH, n=24), mCherry-DHPH (DHPH, n=26), or Lck-mCherry-DHPH (Lck-DHPH, n=22). For **F)** with either C1-mCherry (Control, n=26), mCherry-PLEKHG2 (FL, n=19), mCherry-DH (DH, n=13), Lck-mCherry-DH (Lck-DH, n=33), mCherry-DHPH (DHPH, n=19), or Lck-mCherry-DHPH (Lck-DHPH, n=35). For **G)** with either C1-mCherry (Control, n=25), mCherry-PLEKHG4 (FL, n=19), mCherry-DH (DH, n=22), Lck-mCherry-DH (Lck-DH, n=26), mCherry-DHPH (DHPH, n=21), or Lck-mCherry-DHPH (Lck-DHPH, n=23). And for **H)** with either C1-mCherry (Control, n=22), mCherry-pRex1 (FL, n=45), mCherry-DH (DH, n=23), Lck-mCherry-DH (Lck-DH, n=19), mCherry-DHPH (DHPH, n=13), or Lck-mCherry-DHPH (Lck-DHPH, n=17)

